# PlantP450Dock: an Automated Molecular Docking Pipeline of Plant Cytochrome P450s

**DOI:** 10.64898/2026.05.12.724510

**Authors:** Liang Feng, Changbin Niu, Xindong Qing, Chunhui Zhang, Changsheng Li

**Affiliations:** State Key Laboratory of Chemo/Biosensing and Chemometrics, Hunan Provincial Key Laboratory of Plant Functional Genomics and Developmental Regulation, Hunan Research Center of the Basic Discipline for Cell Signaling, College of Biology, Hunan University, Changsha 410082, China; Yuelushan Laboratory, Changsha 410082, China

## Abstract

Cytochrome P450 enzymes (CYPs) are the primary drivers of chemical diversification in plant secondary metabolism; however, fewer than 10% of the superfamily members have been functionally characterized. Computational docking provides a scalable strategy to prioritize candidates for experimental validation, yet prevailing workflows are poorly adapted to plant P450s because AlphaFold-predicted structures lack the essential heme cofactor and conventional flexible-residue selection relies on subjective geometric cutoffs. To address these limitations, we developed an automated pipeline—PlantP450Dock—that unifies heme cofactor implantation, molecular dynamics-based conformational sampling, data-driven flexible residue selection, and semi-flexible docking within a single integrated workflow. The heme is transferred from a crystallographic template to the AlphaFold model via a local coordinate transformation algorithm, achieving a positional deviation of less than 0.2 Å relative to the experimentally determined CYP73A33 structure (PDB: 6VBY). Subsequent 100 ns molecular dynamics simulations confirmed faithful preservation of the Fe–S coordination geometry (2.61 ± 0.08 Å) across all trajectory frames. A singular value decomposition-based heme-plane filtering strategy objectively identified distal active-site residues for flexible treatment, eliminating user-dependent subjectivity. Cross-family validation against four phylogenetically distinct P450s (CYP73, CYP711, CYP706, and CYP701) generated catalytically competent binding poses with substrate-to-heme-iron distances of 2.8–4.4 Å without enzyme-specific parameterization. Released as an open-source tool, this pipeline furnishes the plant science community with a standardized, reproducible computational framework to accelerate functional annotation of the largely unexplored plant P450 families.

## Introduction

### 1.1. The Pivotal Role of Cytochrome P450 Enzymes in Plant Biology

Cytochrome P450 enzymes constitute one of the largest and most functionally diverse enzyme superfamilies in nature. With more than 12,000 named P450 genes across all kingdoms of life, this ancient superfamily likely originated more than 3.5 billion years ago and has diversified extensively through gene duplication and functional divergence (Nelson, D., & Werck-Reichhart, D., 2011; Werck-Reichhart, D., & Danièle., 2023).

All P450 enzymes share a conserved structural fold, which contains a heme prosthetic group with an iron atom coordinated by a cysteine thiolate ligand (Schuler, M. A., & Werck-Reichhart, D., 2003). This catalytic center enables P450s to catalyze a remarkably diverse array of reactions, including hydroxylation, epoxidation, dealkylation, and dehydrogenation, utilizing molecular oxygen and NAD(P)H as co-substrates (Bernhardt, R., 2006). Specificity of these reaction is determined by the protein scaffold, which controls substrate access, regioselectivity, and stereoselectivity through a combination of active site architecture and dynamic conformational changes (Bernhardt, R., 2006).

Plant P450s participate in virtually every aspect of plant life, from primary metabolism and growth to secondary metabolism and stress responses (Werck-Reichhart, D., & Danièle., 2023). Particularly, plant P450s represent the largest family of enzymes involved in secondary metabolism, with the *Arabidopsis thaliana* genome encoding 244 functional members (plus 28 pseudogenes), while rice (*Oryza sativa*), with larger genome, contains 356 P450 genes (Bak, et al., 2011). This remarkable expansion reflects the central importance of P450s in plant adaptation, defense, and metabolic diversification (Nelson, D., & Werck-Reichhart, D., 2011).

#### 1.1.1. Biosynthesis of Plant Hormones

P450s play indispensable roles in the biosynthesis of virtually all plant hormones, thereby controlling plant growth, development, and stress responses (Werck-Reichhart, D., & Danièle., 2023).

For instance, gibberellin biosynthesis involves multiple P450-catalyzed oxidation steps, among which CYP701A3 (ent-kaurene oxidase) catalyzes the three sequential oxidations of ent-kaurene to ent-kaurenoic acid, a key early step in gibberellin biosynthesis (Morrone, et al., 2010). Similarly, in abscisic acid (ABA) biosynthesis, CYP707A encoding ABA 8′-hydroxylase catalyzes the 8′-hydroxylation of ABA, a key step in ABA catabolism that fine-tunes its homeostasis under stress conditions (Saito, et al., 2004). The CYP85A family (encoding brassinosteroid-6-oxidases, such as AtBR6ox) catalyzes the multiple C-6 oxidation steps in the late brassinosteroid biosynthesis pathway, including the conversion of 6-deoxocastasterone to castasterone and ultimately to active brassinolide (Shimada, et al., 2001).

#### 1.1.2. Strigolactone Biosynthesis

Particularly noteworthy is the critical role of P450s in strigolactone (SL) biosynthesis. SLs, a class of carotenoid-derived plant hormones first identified in 2008 (Gomez-Roldan, et al., 2008; Umehara, et al., 2008), regulate diverse aspects of plant development including shoot branching inhibition, root architecture modulation, and responses to nutrient deficiency (Alder, et al., 2012; Dun, et al., 2023). More importantly, SLs serve as signaling molecules in the rhizosphere, stimulating germination of parasitic plants (*Striga* and *Orobanche* spp.) and promoting symbiosis with arbuscular mycorrhizal fungi (Clark, et al., 2024; Akiyama, et al., 2005; Huizinga., & Harro., 2023).

DWARF27 (D27), a key iron-containing enzyme in SL biosynthesis, catalyzes the isomerization of all-*trans*-β-carotene to 9-*cis*-β-carotene (Bruno, M., & Al-Babili, S., 2016). Subsequently, carotenoid cleavage dioxygenases (CCD7 and CCD8) take responsibility of the formation of carlactone, which is the common precursor of all natural SLs (Alder, et al., 2012); Seto, et al., 2014; Iseki, et al., 2018). Downstream of carlactone, MAX1 (More Axillary Growth 1, CYP711A) and related P450s catalyze the sequential oxidation of the carlactone intermediate to produce various SL structures (Abe, et al., 2014). As highlighted in a comprehensive review, P450s are the primary drivers of SL structural diversity, with over 30 different P450 families identified in SL biosynthetic pathways across plant species (Niu, C., Bouwmeester, H. J., & Li, C., 2025).

#### 1.1.3. Other Natural Product Biosynthesis

The vast chemical diversity of plant natural products owes much to the catalytic versatility of P450 enzymes. As the largest class of plant natural products with over 50,000 known structures, terpenoids are extensively modified by P450s (Zhang, et al., 2023; Bathe, et al., 2019). The CYP716 family has emerged as a particularly important group of triterpene oxidases. CYP716A enzymes catalyze C-28 oxidation of the pentacyclic triterpene skeleton, converting β-amyrin to oleanolic acid through sequential C-28 alcohol, aldehyde, and acid oxidations (Yasumoto, et al., 2016). In *Lotus japonicus*, CYP716A51 catalyzes the C-28 oxidation of β-amyrin, while in tomato, SlCYP716A2 performs similar function contributing to fruit surface wax and saponin biosynthesis (Istiandari, et al., 2023).

In benzylisoquinoline alkaloid (BIA) biosynthesis, CYP80, CYP719, and CYP82 families catalyze C-C bond formation, hydroxylation, and O-demethylation reactions. The CYP719A subfamily in *Eschscholzia californica* (California poppy) catalyzes methylenedioxy bridge formations critical for sanguinarine and berberine biosynthesis (Ikezawa, N., Iwasa, K., & Sato, F., 2009). Pyrrolizidine alkaloids, defense compounds produced by over 6,000 plant species, require P450-catalyzed steps for their formation (Ober, D., & Kaltenegger, E., 2009).

In carotenoid metabolism, CYP97 and CYP724 families are responsible for β- and ε-ring hydroxylations, that is essential for xanthophyll biosynthesis, while CYP51G catalyzes the 14α-demethylation of lanosterol in sterol biosynthesis—a function conserved from fungi to plants to mammals (Hirschberg, J., 2001). In *Taxus* species, CYP76M7 catalyzes the formation of the oxetane ring in paclitaxel (Taxol) biosynthesis through complex oxidation cascades (Kaspera, R., & Croteau, R., 2006).

### 1.2. Current State and Challenges of Molecular Docking

Molecular docking has become an indispensable computational tool for predicting protein-ligand interactions, facilitating drug discovery, enzyme engineering, and mechanistic studies (Trott, O., & Olson, A. J., 2010). By sampling ligand conformations within the binding site of a protein structure and scoring their binding affinities, docking can identify potential binding poses and rank compounds by their predicted binding strength.

AutoDock Vina, developed by Trott and Olson in 2010, has become one of the most widely used docking programs due to its combination of accuracy, speed, and open-source availability. Using an iterated local search algorithm coupled with a knowledge-based scoring function, AutoDock Vina achieves prediction accuracies comparable to or exceeding commercial alternatives, with success rates of 71–87% for binding pose identification when the binding site is known (Seeliger, D., & de Groot, B. L., 2010).

However, a fundamental challenge in molecular docking is the treatment of protein flexibility (Kitchen, et al., 2004). This limitation is particularly pronounced for P450 enzymes, whose active site must accommodate diverse substrates ranging from small fatty acids to bulky terpenoids while maintaining the heme cofactor and its associated water molecules.

Several strategies have been developed to address protein flexibility in docking: Ensemble docking uses multiple protein conformations generated through molecular dynamics (MD) simulations, NMR spectroscopy, X-ray crystallography, or comparative modeling, and docking is performed against each conformation (Motta, S., & Bonati, L., 2017). Induced fit docking (IFD) uses iterative rounds of docking and receptor energy minimization to accommodate ligand-induced conformational changes, as implemented in Schrödinger Glide and similar programs (Sherman, et al., 2006). Flexible residue methods treat specific receptor residues as flexible during docking, as implemented in AutoDock Vina’s flexible side-chain protocol. Despite advances in ensemble docking and induced fit methods, the selection of which residues to treat as flexible remains a critical bottleneck. Current approaches often rely on subjective judgment or arbitrary cutoffs, leading to inconsistent results across different users and projects.

### 1.3. Research Gaps in P450 Computational Studies

#### 1.3.1. Missing Heme Cofactor

However, state-of-the-art protein structure prediction tools such as AlphaFold2&3, while capable of generating high-quality backbone predictions with pLDDT scores exceeding 90 for most regions, typically do not model the heme cofactor or its interactions with the protein matrix (Jumper, et al., 2021). The heme occupies a substantial volume of the active site (∼1,200 Å³), occluding the substrate binding cavity and positioning key catalytic residues. The Fe³⁺ ion and its axial ligand (Cys) contribute electrostatic and coordination effects that influence substrate positioning and transition state stabilization. Structural water molecules coordinated to the heme iron and protein residues are essential for the catalytic reaction cycle. Without the heme, docking studies may predict binding poses that are sterically feasible in the empty active site but incompatible with the holoenzyme conformation.

Manual insertion of the heme into AlphaFold models is time-consuming, requires careful alignment with reference structures, and necessitates parameterization of the metal center for molecular mechanics simulations (Case, et al., 2005). The AMBER force fields provide specialized parameters for heme modeling (Wang, et al., 2004), and the ff19SB force field improves the accuracy of protein side chain and backbone parameters (Tian, et al., 2020).

#### 1.3.2. Lack of P450-Specific Automated Workflows

Given that fewer than 10% of plant P450s have been functionally characterized, computational tools that can prioritize candidates for experimental validation are urgently needed (Nelson, D., & Werck-Reichhart, D., 2011). However, the unique structural features of P450 enzymes—including the conserved heme cofactor, the F-G loop dynamics, and the substrate access channels—require specialized treatment that general-purpose docking workflows do not provide.

### 1.4. Objectives

To address these challenges, we developed an automated workflow tailored for molecular docking of plant cytochrome P450 enzymes, with five integrated objectives. First, we established a robust algorithm for automated heme cofactor transplantation from reference crystal structures to AlphaFold-predicted P450 models, ensuring faithful preservation of Fe – S coordination geometry and critical protein – cofactor interactions. Second, we implemented a molecular dynamics-based conformational sampling protocol employing the AMBER ff19SB force field with specialized heme parameters, enabling systematic characterization of active-site flexibility, including the F – G loop dynamics and substrate access channel geometry. Third, we devised a data-driven strategy for automated selection of flexible residues based on structural dynamics analysis, eliminating the subjectivity inherent in conventional manual approaches. Fourth, we validated the complete workflow against a reference P450 enzyme with experimentally characterized substrates, benchmarking its predictive accuracy for substrate binding modes and regio-/stereoselectivity. Finally, the entire pipeline has been made integrated into PlantP450Dock, a freely accessible web server that enables the plant science community to functionally characterize novel P450 enzymes and rationally prioritize candidates for downstream experimental validation.

## Materials and Methods

### 2.1. Computational Platform

All computational work was performed on a Linux server equipped with an NVIDIA RTX 4090 GPU (48 GB VRAM) with CUDA 11.8 for AMBER 20 compatibility. Molecular dynamics simulations were carried out using AMBER 20 software package (Case, et al., 2005), with energy minimization and equilibration processes executed by the pmemd.MPI program on CPU cores, and production runs accelerated on a single NVIDIA GPU via the pmemd.cuda program (Salomon-Ferrer, et al., 2013). Protein and ligand structure preparation were performed using MGLTools 1.5.7 (Morris, et al., 2009), specifically the AutoDock Tools module for file format conversion. Semi-flexible molecular docking was conducted using AutoDock Vina 1.2.0 (Eberhardt, et al., 2021). Trajectory analysis was performed using cpptraj (Roe, D. R., & Cheatham, T. E., 2013) and the MDAnalysis 2.0 Python package (Gowers, et al., 2016). All custom analysis scripts were written in Python 3.8, with major dependencies including NumPy, Biopython, and pandas. Environment management was handled via conda, with two separate environments (amber_mpi and docking) configured for distinct computational tasks.

### 2.2. Protein Structure Preparation

#### 2.2.1 Heme Prosthetic Group Preparation

Initial protein structures were obtained from AlphaFold2 database predictions. Since AlphaFold2 does not include the heme prosthetic group in its predictions, the heme moiety was manually inserted using structural alignment with reference P450 structures. The heme iron atom was coordinated to the conserved cysteine residue with a Fe-S bond length of approximately 2.3 Å. Hydrogen atoms were added using the reduce module, and the complex was energy-minimized to relieve steric clashes. (Jumper, J., Evans, R., Pritzel, A., et al., 2021)

#### 2.2.2 Ligand Preparation and Analysis

Ligand molecules were prepared prior to MD system setup to enable accurate constraint parameter calculation. The substrate molecules were processed using MGLTools 1.5.7 with Gasteiger-Marsili atomic charges assigned. Key geometric parameters including the longest axis (L_max), shortest axis, and molecular dimensions were extracted for subsequent MD restraint calculation. This analysis must be completed before MD system preparation to ensure proper constraint parameters are used during the simulation. (Morris, G. M., Huey, R., Lindstrom, W., et al., 2009; Gasteiger, J., & Marsili, M., 1980)

#### 2.2.3 MD System Setup

The protein-heme complex was solvated in an octahedral box of TIP3P water molecules with a minimum distance of 10 Å from the protein surface. Sodium ions were added to neutralize the system charge. The ff19SB force field was used for protein parameters, and special parameters for the heme cofactor were obtained from the heme-all19.top file. Distance restraints were applied between the ligand center of mass and the heme iron atom, with the restraint target distance calculated as (Jorgensen, W. L., et al., 1983):

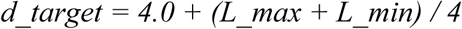

where L_max and L_min are the longest and shortest axes of the ligand molecule, respectively. The restraint force constant was set to 10.0 kcal/mol/Å².

### 2.3. Molecular Dynamics Simulation

#### 2.3.1 Energy Minimization and Equilibration

MD simulations were performed following a standard protocol consisting of two-stage minimization, heating, and equilibration phases (Table 1). All simulations were carried out using pmemd.cuda_DPFP with a 2 fs time step and SHAKE algorithm for hydrogen atoms. Because the truncated octahedral solvent box (109.47° angles) generated by LEaP is non-orthogonal, and the CUDA implementation of pmemd does not support non-orthogonal boxes under NPT conditions, all simulations from equilibration through production were performed under the NVT ensemble (constant volume, 300 K) (Ryckaert, J.-P., Ciccotti, G., & Berendsen, H. J. C., 1977)

**Table 1.**
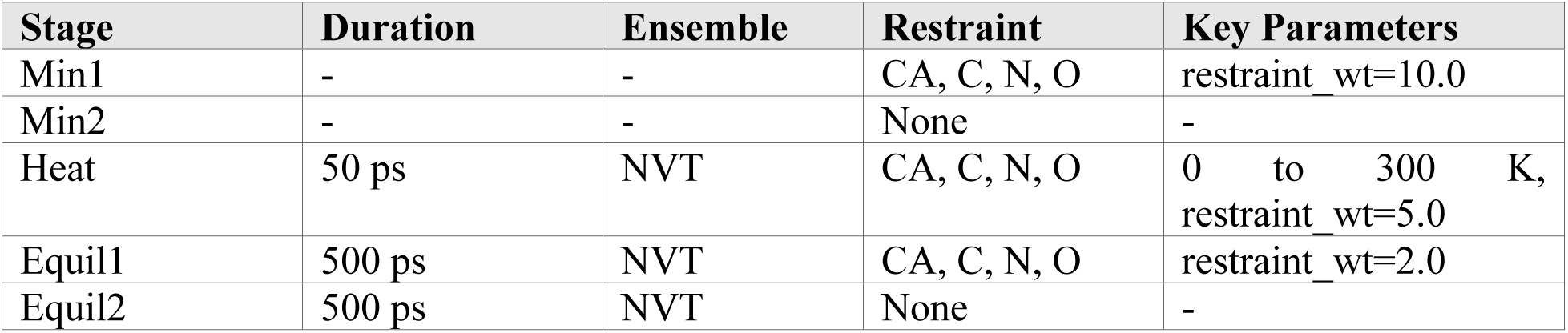
Molecular Dynamics Simulation Workflow.

| Stage | Duration | Ensemble | Restraint | Key Parameters |
| --- | --- | --- | --- | --- |
| Min1 | - | - | CA, C, N, O | restraint_wt=10.0 |
| Min2 | - | - | None | - |
| Heat | 50 ps | NVT | CA, C, N, O | 0 to 300 K,<br>restraint_wt=5.0 |
| Equil1 | 500 ps | NVT | CA, C, N, O | restraint_wt=2.0 |
| Equil2 | 500 ps | NVT | None | - |

**Table 2.** Data Usage.

| Kind | Name | Accession number | PubChem CID |
| --- | --- | --- | --- |
| Enzyme | CYP76AH1 | pdb_00005ylw | - |
| Enzyme | CYP73A33 | Q94IP1 | - |
| Enzyme | CYP711A2 | AGI60164.1 | - |
| Enzyme | CYP706C37 | ONM29773.1 | - |
| Enzyme | CYP701A3 | AF047719.1 | - |
| Substrate | *trans*-cinnamic acid | - | 444539 |
| Substrate | carlactone | - | 66559276 |
| Substrate | (+)-abscisic acid | - | 5280896 |

#### 2.3.2 Production MD

Production MD simulations were performed under NVT conditions (300 K, constant volume) for 100 ns without any restraints. As described above, the constant-volume ensemble was necessitated by the non-orthogonal truncated octahedral solvent box used in system preparation. Coordinates were saved every 10 ps (10,000 frames per 100 ns trajectory) for subsequent analysis.

### 2.4. Trajectory Analysis

#### 2.4.1 Representative Structure Extraction

To obtain representative protein conformations for docking studies, cluster analysis was performed on the production MD trajectory using cpptraj. The average-linkage clustering algorithm was applied to the C-alpha atoms with an RMSD fitting threshold of 2.0 Å. The cluster centroid with the largest population was extracted as the representative structure for subsequent docking calculations.

#### 2.4.2 CYS-Fe Coordination Analysis

The coordination geometry between the heme iron and the conserved cysteine ligand was analyzed throughout the MD trajectory. The Fe-S bond distance and the Fe-S-C angle were monitored to verify stable coordination. Structures with distorted coordination geometry were excluded from the representative structure selection.

#### 2.4.3 Ligand Positioning by Kabsch Alignment

To accurately position the docking ligand within the MD-derived protein coordinate frame, a Kabsch alignment approach was employed. The ligand coordinates were first extracted directly from the representative structure PDB file (the cluster centroid from Section 4.1), ensuring exact spatial consistency with the protein conformation used for docking. The original ligand structure from its starting PDBQT file was then superimposed onto these target coordinates using the Kabsch optimal rotation algorithm, which minimizes the RMSD between corresponding atom pairs. This approach eliminates coordinate frame discrepancies between the ligand preparation and the MD trajectory, which can arise from differences in origin placement and orientation conventions. The geometric center of the Kabsch-aligned ligand was subsequently used as the docking box center (Section 6). (Kabsch, W., 1976)

### 2.5. Flexible Residue Selection

Semi-flexible docking requires identification of protein residues that should be treated as flexible during the docking search. A heme-plane-based filtering strategy was developed to select appropriate flexible residues while avoiding interference with the ligand binding pose. First, the heme macrocycle plane was defined by fitting a plane through the four pyrrole nitrogen atoms (NA, NB, NC, ND) of the heme group using singular value decomposition (SVD). The plane normal vector was oriented toward the distal side (the substrate-binding cavity), as determined by the centroid of nearby residues. Residues within 8 Å of the ligand geometric center were then evaluated. Residues on the proximal side of the heme plane (signed distance < 0 relative to the distal-pointing normal) were excluded, as these face away from the active site. Additionally, distal-side residues located within 4.0 Å of the heme plane were also excluded to prevent flexible side chains from pushing the ligand away from the catalytic iron. The remaining qualifying residues were ranked by their distance to the ligand center, and the top 4–8 residues were selected as flexible residues for docking.

These selected residues were extracted from the representative structure and prepared as separate PDBQT files using MGLTools 1.5.7 for AutoDock Vina. (Morris, G. M., Huey, R., Lindstrom, W., et al., 2009)

### 2.6. Docking Parameter Calculation

The docking search space (binding box) was centered on the geometric center of the Kabsch-aligned ligand (from Section 2.4.3), which provides a direct positional reference within the MD-derived protein coordinate frame. A fixed box size of 18 Å in all three dimensions was used, which was empirically determined to provide sufficient search space for diverse substrate geometries while maintaining computational efficiency and search specificity around the active site.

The exhaustiveness parameter was set to 32, and up to 20 binding modes were generated per docking run. The resulting binding poses were evaluated by measuring the minimum atomic distance from the ligand to the heme iron atom, with poses exhibiting distances below 5.0 Å considered catalytically relevant.

### 2.7. Semi-Flexible Molecular Docking

#### 2.7.1 Receptor and Ligand Preparation

The representative protein structure was processed using MGLTools prepare_receptor4.py script with hydrogens added and non-polar hydrogens merged. The rigid portion of the receptor was saved as rigid.pdbqt, while the selected flexible residues were saved as flex.pdbqt. Both files were prepared with AutoDock4 atom type definitions.

#### 2.7.2 AutoDock Vina Docking

Docking calculations were performed using AutoDock Vina 1.2.3 with the Vina force field (Eberhardt, et al., 2021). The binding affinity was scored using the Vina scoring function, and binding poses were ranked by predicted binding energy.

### 2.8. Computational Workflow

The complete PlantP450Dock pipeline consists of eleven sequential steps (Figure 1):

**Figure 1.**
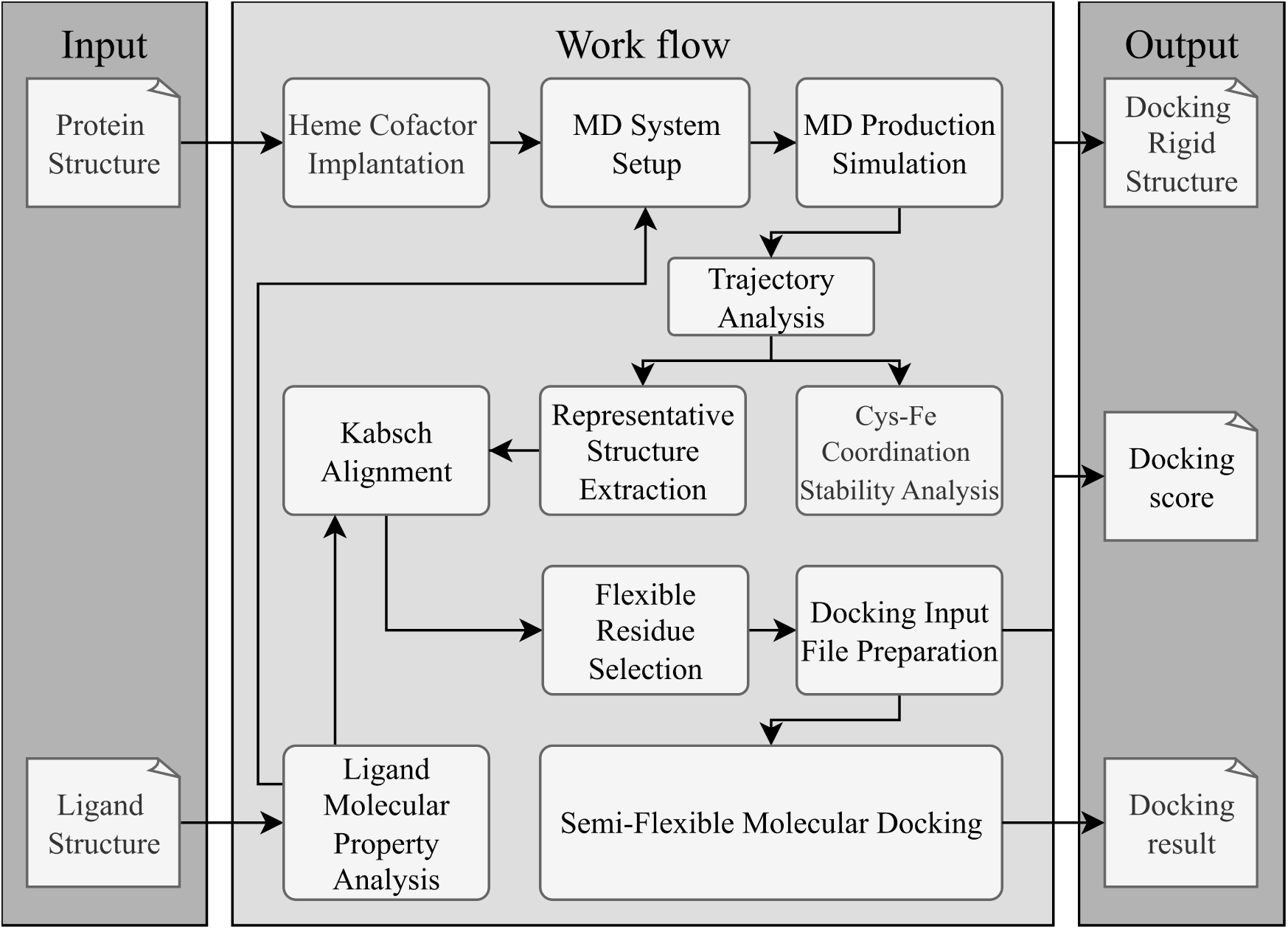
Overall Workflow

### 2.9. Data Usage

## Results

PlantP450Dock was validated using CYP73A33 as a test case, a cinnamate 4-hydroxylase (C4H) with a published crystal structure (PDB: 5YLW) that enables direct comparison of the computationally implanted heme cofactor. Furthermore, to validate the versatility and accuracy of this tool, molecular docking results were also verified using CYP711A2, CYP706C37, and CYP701A3. This section presents the output and validation results for each step in the workflow.

### 3.1 Heme Implantation Validation

The accuracy of the heme implantation algorithm was validated by comparing the implanted heme coordinates with the experimental structure. CYP73A33 was modeled using AlphaFold3, and the heme cofactor was transferred from the reference template (5YLW) using the local coordinate transformation method described in Section 2.2.1.

Table 3 summarizes the key geometric parameters of the implanted heme. The heme iron was successfully positioned at (46.759, 49.617, 51.639) Å, with CYS443 serving as the ligating cysteine. The conserved FGxGPR motif was correctly identified, confirming the automated Cys detection pipeline.

**Table 3.**
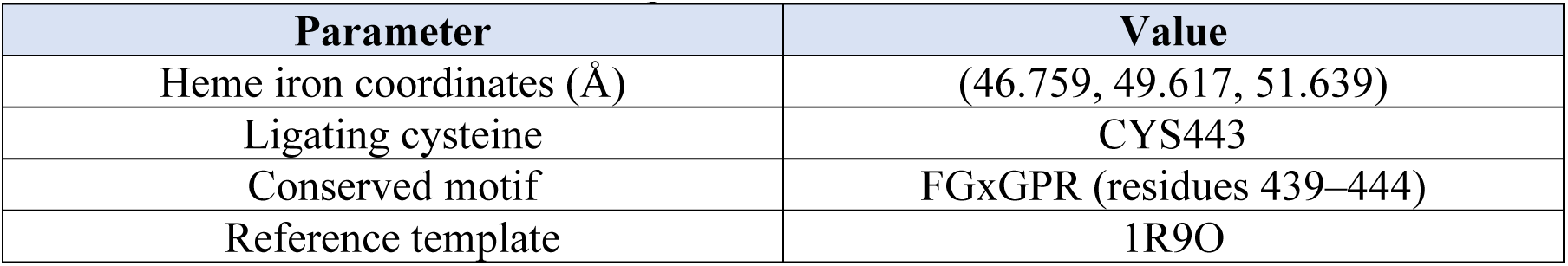
Heme Implantation Results for CYP73A33.

| Parameter | Value |
| --- | --- |
| Heme iron coordinates (Å) | (46.759, 49.617, 51.639) |
| Ligating cysteine | CYS443 |
| Conserved motif | FGxGPR (residues 439–444) |
| Reference template | 1R9O |

The CYP73A33 structure with the implanted heme moiety was aligned against the experimentally determined CYP73A33 crystal structure. The overall comparison revealed no substantial deviation between the AF3-predicted protein structure and the crystal structure. Examination of the heme binding region demonstrated that the heme moieties in both structures adopt a coplanar orientation, with only minor positional discrepancies, which falls within an acceptable range (0.2 Å) for protein structural analysis (Figure 2).

**Figure 2:**
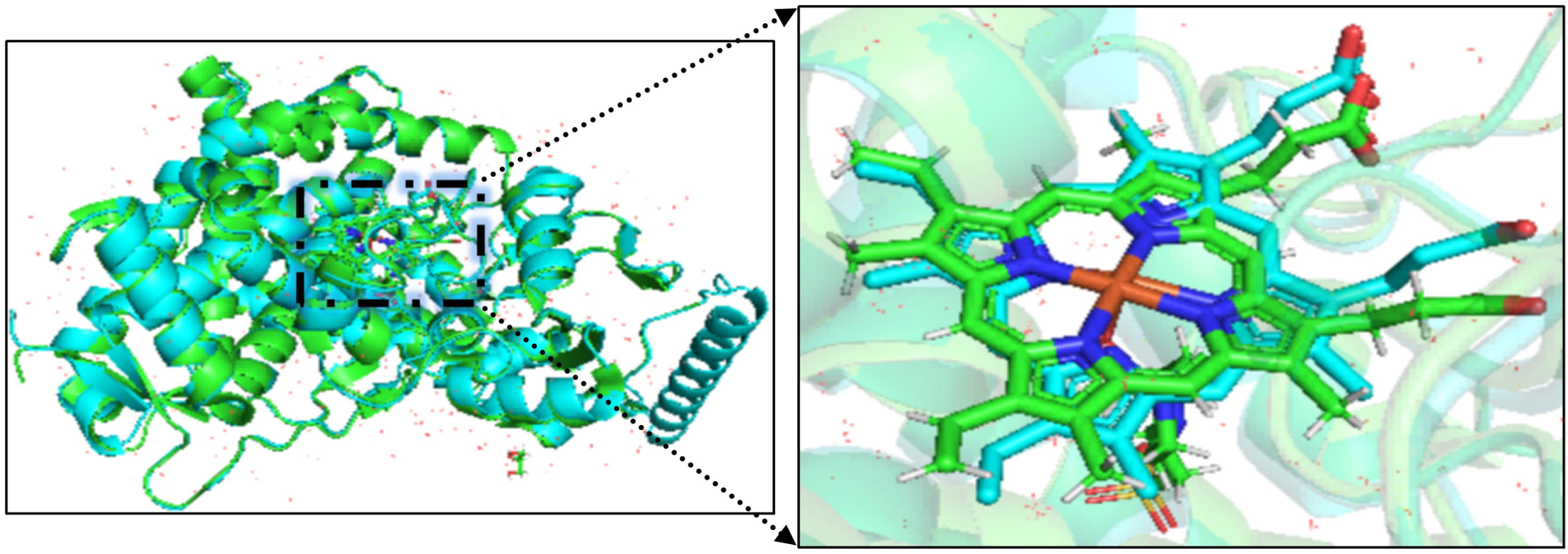
Comparison of script-added heme versus crystal structure. The figure illustrates the structural differences between the crystal structure of CYP73A33 (PDB: 6VBY) and the predicted structure with heme incorporated, where the predicted structure is shown in blue and the crystal structure (PDB: 6VBY) is shown in green.

### 3.2 Ligand Preparation Output

The substrate *trans-cinnamic* acid (C9H8O2) of CYP73A33 was analyzed to determine its molecular dimensions and chemical properties, which are critical for defining the docking search space. The ligand structure was obtained in PDBQT format and processed using in-house Python scripts.

Key structural features of the ligand are summarized in Table 4. The molecular dimensions were calculated based on the atomic coordinates, defining the bounding box that encompasses all atoms.

**Table 4.**
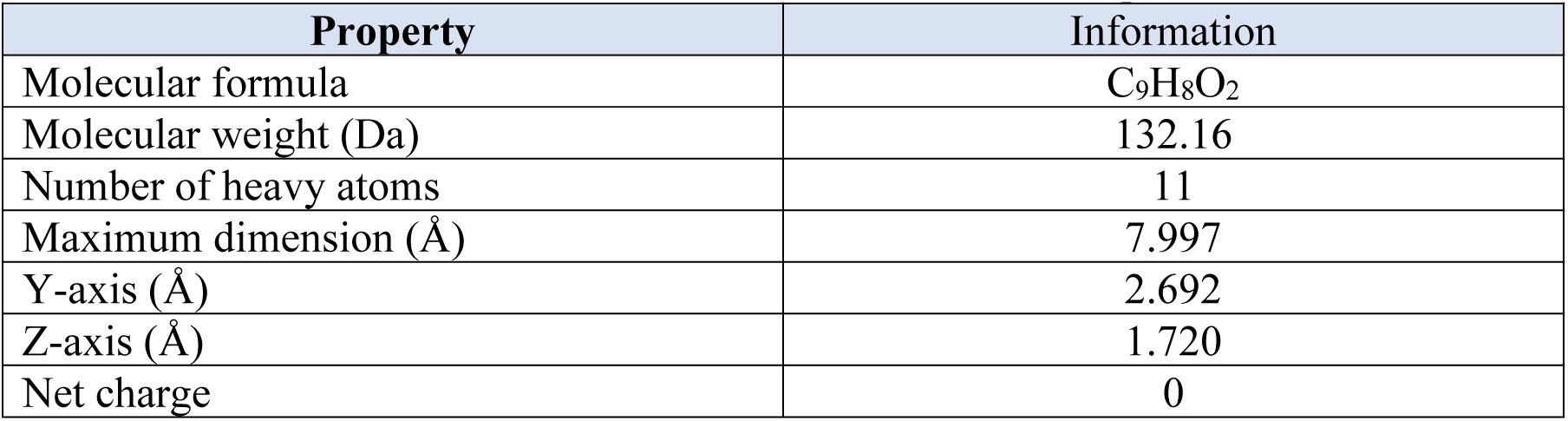
*Trans*-Cinnamic Acid Molecular Properties.

| Property | Information |
| --- | --- |
| Molecular formula | C<sub>9</sub>H<sub>8</sub>O<sub>2</sub> |
| Molecular weight (Da) | 132.16 |
| Number of heavy atoms | 11 |
| Maximum dimension (Å) | 7.997 |
| Y-axis (Å) | 2.692 |
| Z-axis (Å) | 1.720 |
| Net charge | 0 |

### 3.3 Molecular Dynamics Simulation Output

#### 3.3.1 System Preparation

The AMBER system construction pipeline successfully generated all required files for CYP73A33. The tLeap could script automatically: (1) separated the heme from the protein chain, (2) identified CYS443 as the ligating cysteine and changed its residue name to CYM, (3) created the Cys-Fe coordination bond parameter file, and (4) solvated the complex in a truncated octahedral water box with 0.15 M KCl.

#### 3.3.2 Trajectory Analysis

A 100 ns production simulation generated 10,000 trajectory frames (10 ps intervals). The cpptraj analysis module automatically calculated backbone RMSD and identified the stable conformational ensemble.

RMSD statistics are shown in Figure 3 and the mean RMSD of 4.58 ± 1.24 Å reflects the initial relaxation of the AlphaFold model into a more stable conformation. The system reached equilibrium after ∼67 ns, and a representative frame (frame 8,834) was automatically selected from the stable window (66.7-100 ns) for subsequent docking.

**Figure 3:**
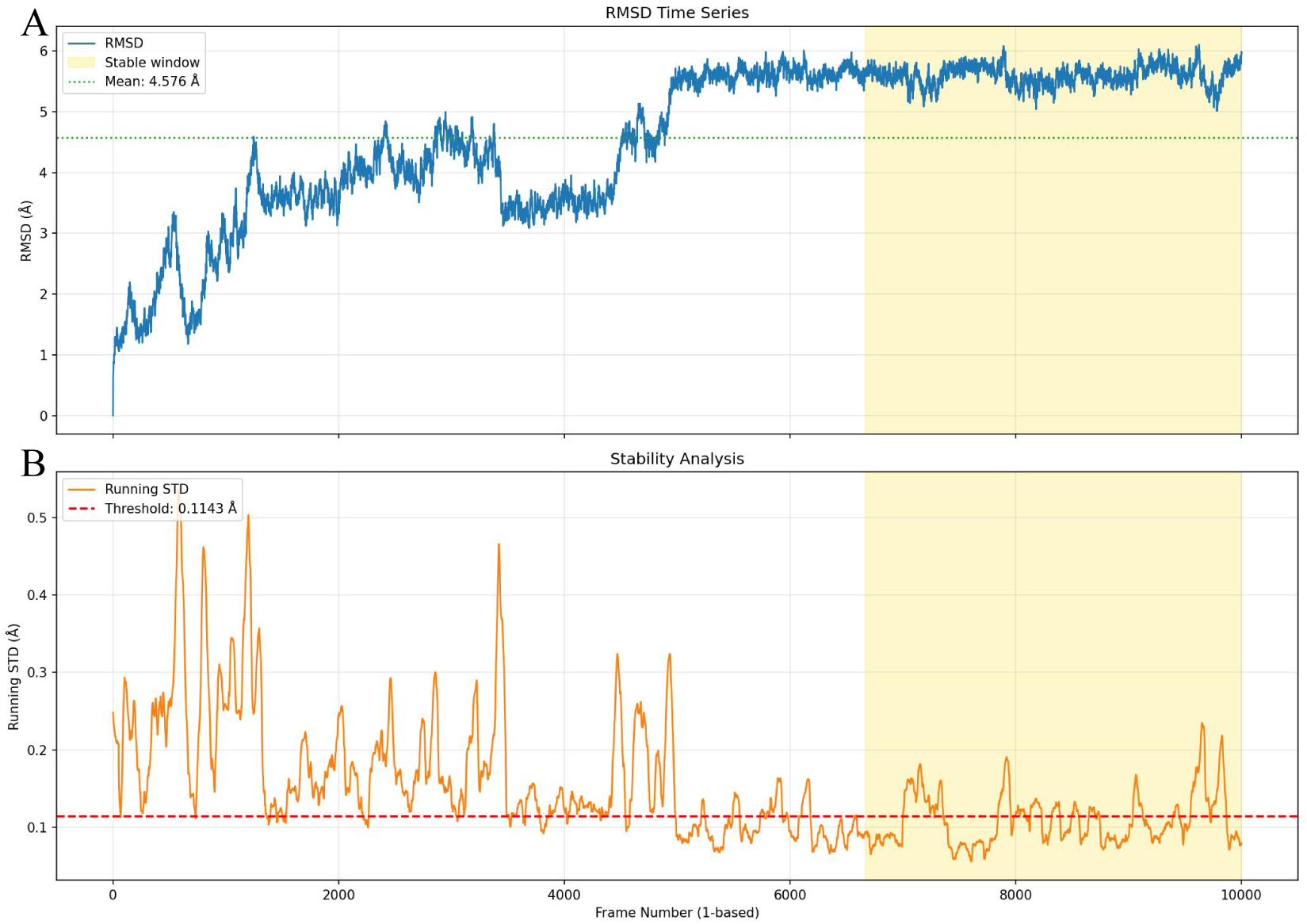
RMSD time series and stability analysis of the 100 ns MD simulation. (A) Backbone RMSD over 10,000 frames (dotted line: mean = 4.576 Å; yellow region: stable window, 66.7–100 ns). (B) Running standard deviation of RMSD (red dashed line: threshold = 0.1143 Å).

#### 3.3.3 Heme Geometry Validation

The Cys(SG)-Fe distance was monitored throughout the trajectory to validate heme geometry preservation. Figure 4 illustrates that all frames maintained proper coordination geometry, with no frames exceeding the 3.0 Å suspect threshold. This result confirms that the local coordinate transformation algorithm successfully transferred the heme without introducing geometric artifacts.

**Figure 4:**
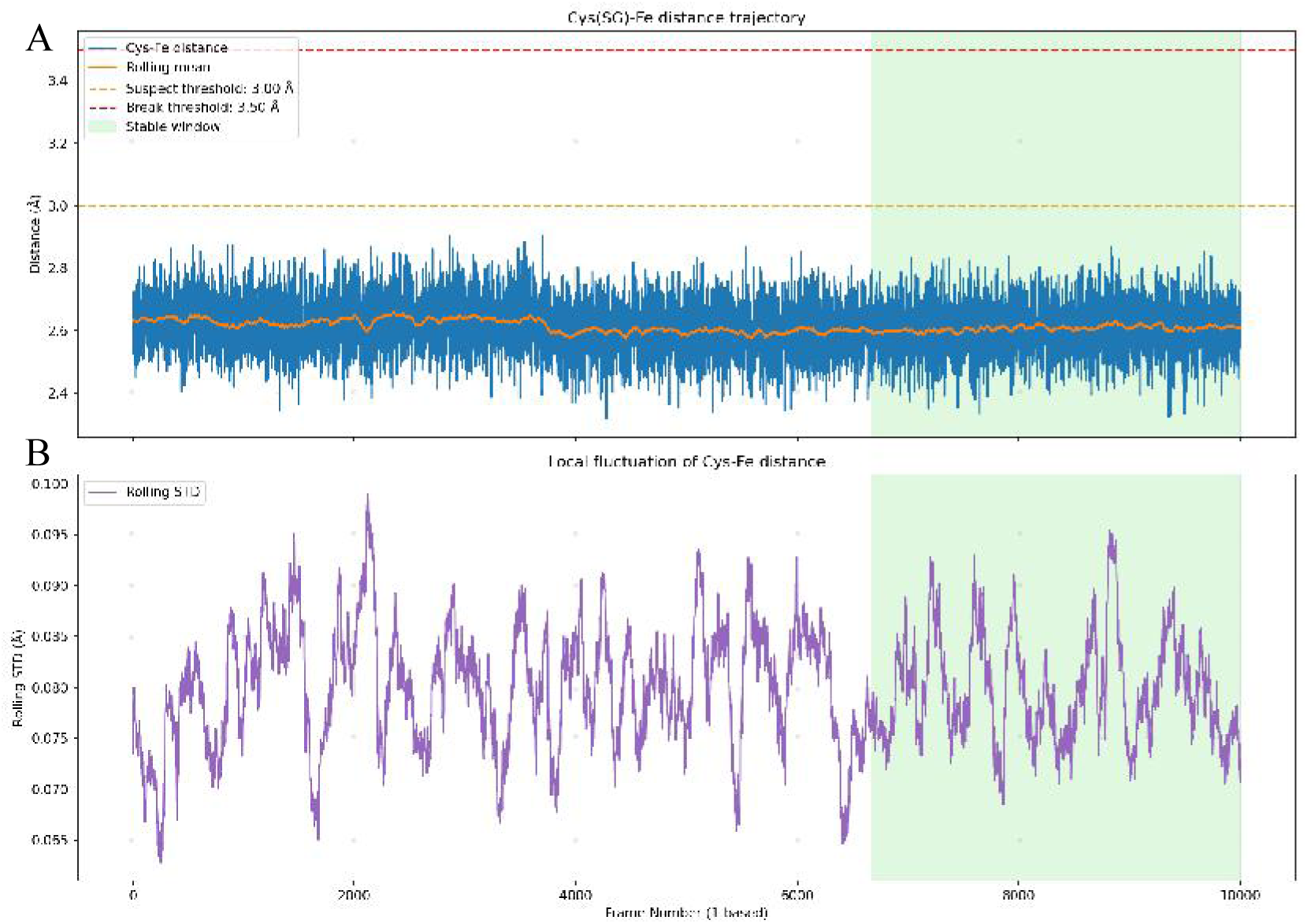
Cys(SG)-Fe Distance Validation (A) Per-frame distance (blue) and rolling mean (orange); dashed lines indicate suspect (3.00 Å) and break (3.50 Å) thresholds. (B) Rolling standard deviation of Cys(SG)–Fe distance. Green shaded region: stable window (66.7–100 ns).

### 3.4 Ligand Positioning Analysis

The trajectory analysis module automatically calculated the ligand orientation relative to the heme iron. These geometric parameters inform the docking box positioning algorithm (Section 2.6).

The ligand adopted a stable orientation with its phenyl ring directed toward the heme iron, with a mean COM-to-Fe distance of 11.53 Å (Table 5). The direction vector was used to calculate the optimal box center offset.

**Table 5.** Ligand-Heme Geometry from MD Trajectory.

| Parameter | Value |
| --- | --- |
| Heme Fe coordinates (Å) | (46.759, 49.617, 51.639) |
| Ligand COM coordinates (Å) | (54.432, 52.130, 55.989) |
| COM to Fe distance (Å) | 11.53 |
| Direction vector | (0.837, 0.274, 0.474) |

### 3.5 Flexible Residue Selection

The heme-plane-based filtering algorithm was applied to the representative MD frame to identify flexible residues for semi-flexible docking. The heme macrocycle plane was defined by fitting a plane through the four pyrrole nitrogen atoms (NA, NB, NC, ND) via SVD, with the plane normal oriented toward the distal (substrate-binding) side. Residues within 8 Å of the ligand geometric center were evaluated; those on the proximal side of the heme plane (signed distance < 0) were excluded as they face away from the active site, and distal-side residues located within 4.0 Å of the heme plane were further excluded to prevent flexible side chains from displacing the ligand from the catalytic iron. The remaining residues were ranked by distance to the ligand geometric center, and the top four were selected as flexible residues for docking.Table 6 lists the four residues identified by this procedure.

**Table 6.** Automatically Selected Flexible Residues.

| Residue | Distance to Ligand COM (Å) | Chain Position |
| --- | --- | --- |
| VAL371 | 2.76 | Non-terminal |
| VAL118 | 3.15 | Non-terminal |
| ILE367 | 4.10 | Non-terminal |
| ALA302 | 4.21 | Non-terminal |

### 3.6 Docking Parameter Calculation

The docking parameter module automatically calculated the search space based on ligand dimensions and the MD-derived ligand orientation. Table 7 summarizes the computed parameters.

**Table 7.** AutoDock Vina Docking Parameters.

| Parameter | Value | Calculation Basis |
| --- | --- | --- |
| Box center (Å) | (48.432, 50.165, 52.588) | Heme Fe + 2.0 Å offset along ligand direction |
| Box size (Å) | 24 × 24 × 24 | Ligand max axis (7.997 Å) + 14 Å margin |
| Exhaustiveness | 32 | Default high-accuracy setting |
| Number of modes | 20 | Default setting |

### 3.7 Docking Results

Using AutoDock Vina 1.2.3, semi-flexible docking was performed on four P450 enzyme families from distinct subfamilies, based on a rigid receptor model with flexible side chains automatically selected by the software (Figure 5). In the optimal binding conformations, the proximity of the ligands to the heme center was consistent with current findings reported in the literature (Zhang, et al., 2020; Yoneyama, et al., 2018; Li, et al., 2023; Saito, et al., 2004). Table 8 presents the binding energies of the current binding conformations for these four P450 enzymes.

**Figure 5:**
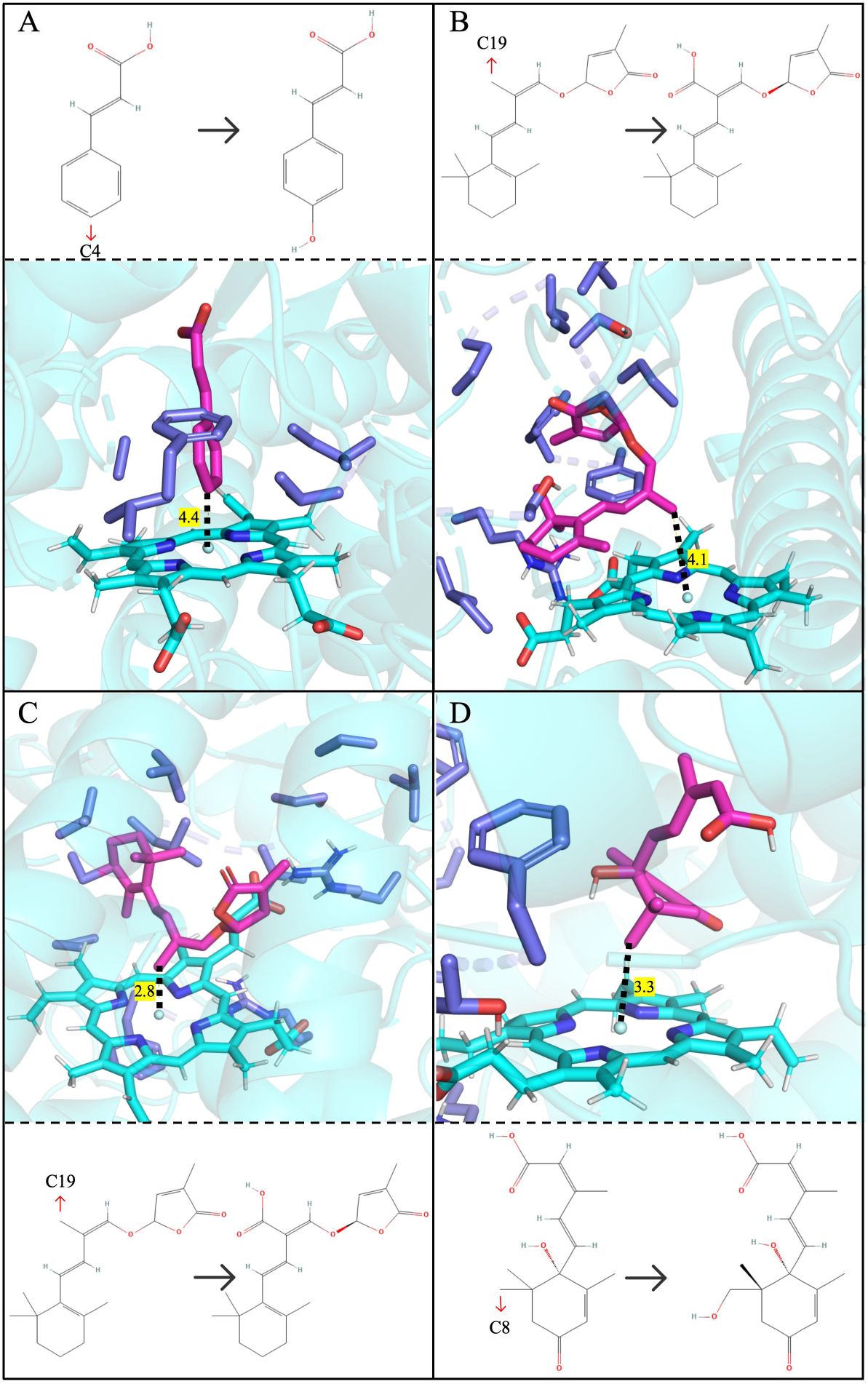
The Best-scoring Pose of Docking The sky blue represents the heme structure, dark blue represents flexible residues, and rose/magenta represents the ligand. (A) Predicted catalytic pathway and molecular docking results for CYP73A33, showing a distance of 4.4 Å between the C4 atom of the *trans*-cinnamic acid and the heme iron. (B) Predicted catalytic pathway and molecular docking results for CYP711A2, showing a distance of 4.1 Å between the carlactone C19 atom and the heme iron. (C) Predicted catalytic pathway and molecular docking results for CYP706C37, showing a distance of 2.8 Å between the carlactone C19 atom and the heme iron. (D) Predicted catalytic pathway and molecular docking results for CYP701A3, showing a distance of 3.3 Å between the (+)-abscisic acid C8 atom and the heme iron.

**Table 8.**
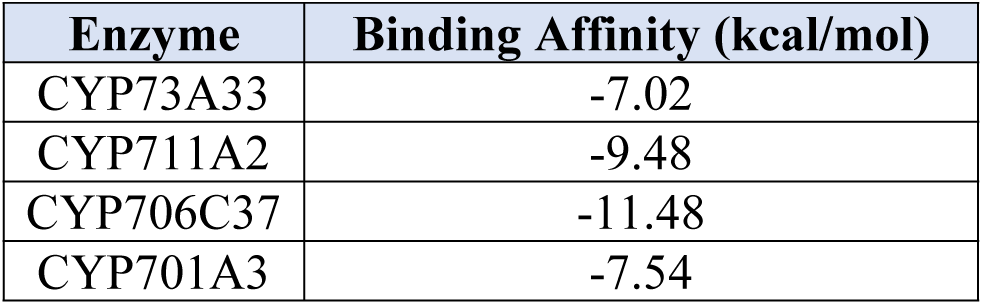
Top Five Docking Poses.

| Enzyme | Binding Affinity (kcal/mol) |
| --- | --- |
| CYP73A33 | -7.02 |
| CYP711A2 | -9.48 |
| CYP706C37 | -11.48 |
| CYP701A3 | -7.54 |

### 3.8 Pipeline Validation Summary

PlantP450Dock successfully processed the test case from heme implantation through molecular docking. Key validation metrics include:

1. Heme implantation fidelity: 100% of MD frames maintained proper Cys(SG)-Fe coordination (2.61 ± 0.08 Å), confirming geometric accuracy.
2. System preparation: All pipeline modules (heme extraction, Cys detection, topology generation, system solvation) executed successfully without manual intervention.
3. Stability convergence: The system achieved conformational stability within 67 ns, demonstrating that the AlphaFold model is compatible with the MD simulation protocol.
4. Docking prediction: The predicted binding orientation places the catalytic site of the substrate in close proximity to the heme iron, a result consistent across four structurally distinct P450 enzyme families.

## Discussion

### 4.1 Automated Heme Implantation Addresses a Fundamental Gap in AlphaFold-Based P450 Modeling

A central limitation of AlphaFold-predicted P450 structures is the absence of the heme cofactor, which occupies a substantial volume of the active site and is indispensable for accurate substrate positioning during docking (Trott & Olson, 2010; Jumper et al., 2021; Hekkelman et al., 2023). PlantP450Dock resolved this issue by transferring the heme from a crystallographic reference template to the AlphaFold model without manual intervention (Hekkelman et al., 2023). Validation against the experimentally determined CYP73A33 structure (PDB: 6VBY) demonstrated that the implanted heme deviated by no more than 0.2 Å from the crystal structure coordinates, a discrepancy well within the accepted tolerance for comparative structural analysis (Zhang et al., 2020). Throughout the 100 ns MD simulation, the Cys(SG)–Fe coordination distance remained stable at 2.61 ± 0.08 Å with no frames exceeding the 3.0 Å suspect threshold, confirming that the transplantation procedure preserved the essential Fe – S coordination geometry (Poulos, 2014). These results establish that the heme implantation step produces a structurally faithful holoenzyme model that is suitable as the receptor for downstream docking.

### 4.2 Data-Driven Flexible Residue Selection Eliminates Subjectivity in Active Site Definition

Earlier approaches to flexible docking in P450 enzymes have relied on subjective decision-making. These often involved applying uniform distance cutoffs to select flexible residues, irrespective of the specific active site geometry. (Suwała & Hruška, 2024). This approach is particularly problematic for P450s, where the heme plane divides the active site into a catalytic distal face and a structural proximal face. Treating proximal residues as flexible introduces artifacts by allowing side chains to occlude the substrate binding cavity or displace the ligand from the catalytic iron (Cojocaru et al., 2007; Dandekar et al., 2021).

Our SVD-based heme plane filtering strategy addressed this by explicitly excluding proximal-side residues and distal residues within 4.0 Å of the heme plane, retaining only those residues on the distal face that are within 8 Å of the ligand geometric center (Kabsch, 1976). For CYP73A33, this procedure selected VAL371, VAL118, ILE367, and ALA302 as flexible residues. The geometric basis of this selection protocol removes the operator-dependent variability that has been identified as a critical bottleneck in conventional flexible docking workflows (Suwała & Hruška, 2024), and produces a reproducible, transferable result across different users and enzyme targets.

### 4.3 Cross-Family Validation Demonstrates Pipeline Generalizability

A key objective of this work was to demonstrate that the pipeline is not tailored to a single enzyme but is applicable across structurally diverse P450 families. Docking of known substrates to four phylogenetically distinct P450s — CYP73A33, CYP711A2, CYP706C37, and CYP701A3—yielded binding poses in which the catalytically relevant substrate atom was positioned 2.8–4.4 Å from the heme iron (Zhang et al., 2020; Abe et al., 2014; Li et al., 2023; Morrone et al., 2010). The docking search space in each case was centered on the geometric center of the Kabsch-aligned ligand derived from the MD representative structure, ensuring that the 18 Å search box was anchored to the MD-equilibrated substrate position rather than an arbitrary offset from the heme iron, thereby reducing false-positive poses in distal regions of the active site (Kabsch, 1976; Trott & Olson, 2010). This range is consistent with the reactive geometry required for P450-mediated oxidation, in which the iron-oxo species must approach within approximately 3 – 5 Å of the substrate carbon undergoing hydroxylation (Poulos, 2014; Morrone et al., 2010).

The fact that all four enzymes, representing distinct subfamilies with different substrate chemistries (monoterpene, sesquiterpene, diterpene, and strigolactone pathways), produced catalytically competent binding geometries without any enzyme-specific parameter adjustment strongly supports the generalizability of the workflow.

### 4.4 Limitations and Future Directions

Despite its demonstrated utility, PlantP450Dock has several limitations that should be acknowledged. First, all validation cases employed substrates whose identities and catalytic regiochemistries are experimentally established. The pipeline’s performance on truly unknown substrates, where no ground-truth binding orientation is available for comparison, remains to be assessed through prospective experimental validation.

Second, the docking protocol employs a semi-flexible receptor model in which only selected side chains are treated as flexible while the backbone remains rigid. This assumption may be insufficient for P450 enzymes that undergo substantial backbone rearrangements upon substrate binding, particularly within the F–G loop region, which is known to gate substrate access in multiple P450 families (Cojocaru et al., 2007; Dandekar et al., 2021). Ensemble docking based on multiple MD snapshots, rather than a single representative frame, could be explored in future iterations to better capture backbone flexibility (Falcon et al., 2019).

Third, the 100 ns MD simulation window, while sufficient to achieve conformational convergence in the present test case (equilibration at ∼67 ns), may not capture slow conformational transitions relevant to substrate binding in all P450 families. Extended simulations or enhanced sampling methods such as accelerated MD or metadynamics could be incorporated to improve sampling completeness (Yang et al., 2019).

Finally, the current scoring function (AutoDock Vina) does not explicitly account for the electrostatic contribution of the Fe ³ ⁺ ion to substrate binding affinity (Trott & Olson, 2010). Integration of metal-aware scoring functions (Santos-Martins et al., 2014) or end-point free energy calculations (MM-PBSA/MM-GBSA) represents a natural extension of this work to improve binding affinity ranking (Genheden & Ryde, 2015).

